# CpG traffic lights are markers of regulatory regions in humans

**DOI:** 10.1101/095968

**Authors:** Abdullah M. Khamis, Anna V. Lioznova, Artem V. Artemov, Vasily Ramensky, Vladimir B. Bajic, Yulia A. Medvedeva

**Affiliations:** King Abdullah University of Science and Technology (KAUST), Computational Bioscience Research Center (CBRC), Computer, Electrical and Mathematical Sciences and Engineering (CEMSE) Division, Thuwal 23955-6900, Saudi Arabia; Institute of Bioengineering, Research Center of Biotechnology, Russian Academy of Sciences, Moscow 119071, Russian Federation; Faculty of Bioengineering and Bioinformatics, Lomonosov Moscow State University, Moscow 119991, Russian Federation; Institute for Information Transmission Problems (Kharkevich Institute), Russian Academy of Sciences, Moscow 127051, Russian Federation; Center for Neurobehavioral Genetics, Semel Institute for Neuroscience and Human Behavior, University of California Los Angeles, Los Angeles, California 90095, USA; Moscow Institute of Physics and Technology, Dolgoprudny, Moscow Region 141701, Russian Federation; Immanuel Kant Baltic Federal University, Kaliningrad 236041, Russian Federation; Vavilov Institute of General Genetics, Russian Academy of Sciences, Moscow 119991, Russian Federation

**Keywords:** Regulation of transcription, DNA methylation, enhancers, CAGE, chromatin states, CpG traffic lights

## Abstract

DNA methylation is involved in regulation of gene expression. Although modern methods profile DNA methylation at single CpG sites, methylation levels are usually averaged over genomic regions in the downstream analyses. In this study we demonstrate that single CpG methylation can serve as a more accurate predictor of gene expression compared to average promoter / gene body methylation. CpG positions with significant correlation between methylation and expression of a gene nearby (named CpG traffic lights) are evolutionary conserved and enriched for exact TSS positions and active enhancers. Among all promoter types, CpG traffic lights are especially enriched in poised promoters. Genes that harbor CpG traffic lights are associated with development and signal transduction. Methylation levels of individual CpG traffic lights vary between cell types dramatically with the increased frequency of intermediate methylation levels, indicating cell population heterogeneity in CpG methylation levels. Being in line with the concept of the inherited stochastic epigenetic variation, methylation of such CpG positions might contribute to transcriptional regulation. Alternatively, one can hypothesize that traffic lights are markers of absent gene expression resulting from inactivation of their regulatory elements. The CpG traffic lights provide a promising insight into mechanisms of enhancer activity and gene regulation linking methylation of single CpG to expression.

## Introduction

Epigenetic regulation of gene expression attracted a lot of research attention over the last decade with cytosine methylation being probably the most well-investigated mechanism. DNA methylation is linked to many normal and pathological biological processes: organism development, cell differentiation, cell identity and pluripotency maintenance (reviewed in [23,42,61]), aging [7], memory formation [15,44], responses to environmental exposures, stress and diet [25,33,45]. There is an increasing evidence of abnormalities in DNA methylation present in various diseases, including metabolic [10], cardiovascular [67], neurodegenerative [53,65] diseases and cancers (reviewed in [4]. For about a decade, DNA demethylating drugs (Decitabine, Azacytidine) are used in clinic for the treatment of acute myeloid leukemia and myelodysplastic syndrome [9]. Recent advances in site-specific editing of DNA methylation [56] suggest the possibility of exploring DNA methylation as a promising target for non-invasive therapies against many diseases linked with aberrant methylation.

Functionally, DNA methylation of the promoter region is tightly associated with the repression of transcription initiation, while methylation of the gene body, on the contrary, increases with the the increased expression intensity (reviewed in [26]). Enhancers, distant regulatory regions, that contribute to the establishment of the correct temporal and cell-type-specific gene expression pattern, have been shown to initiate transcription of short RNAs by PolII [27]. Therefore, it is no surprise that DNA methylation might also regulate the enhancer function as well [19,29,48]. Recent studies support the role of DNA methyltransferase in enhancer-associated transcription [50]. The enhancers locations are more difficult to determine genome-wide than those of genes. Some progress in this direction has been made with the use of histone modifications profiles, transcription factor binding or DNase I hypersensitive sites (DHSs) (reviewed in [57]) or the presence of balanced bidirectional capped transcripts (CAGE) [1]. Yet, due to the difficulties in localization of enhancers, the role of their methylation is not completely clear.

It is important to emphasize that epigenetic profiles vary between cells that belong to the same organism and therefore share the same genetic background. The majority of these epigenetic differences are established during development and can be explained by cell types and tissues in a multicellular organism. Yet, an epigenetic heterogeneity has been observed in the normal tissues of inbred laboratory mice [22] and at the level of single cells [13], suggesting stochasticity in the epigenetic profiles intrinsic to some genome loci but not others [14]. The effect of genome-wide epigenetic stochasticity for gene expression has not been addressed so far in fine details [13].

Contemporary methods to study DNA methylation based on bisulfite sequencing allow detection of single cytosine methylation. Yet, at the step of downstream bioinformatic analysis, methylation levels of several dozens of cytosines are usually averaged with the aim to increase statistical power [5,28]. However, several examples show that changes in methylation of a single CpG affect gene transcription [36]. Recently, we have shown that methylation levels of particular single CpGs are tightly linked to expression for specific cases [40]. We have called such positions CpG traffic lights (CpG TL) and have demonstrated a strong negative selection against them in transcriptional factor binding sites. In this study we show enrichment of CpG TL in transcriptional start sites (TSS), in particular, in poised promoters, enhancers and regions with active chromatin marks, suggesting additional mechanism of transcriptional regulation. Also, a study of methylation at the level of a single CpG dinucleotide allows one to address the issue of methylation heterogeneity. Although allele-specific methylation, being a result of intra individual variations, may contribute to observed methylation heterogeneity, its contribution is reduced when samples from non-related individuals are studied together. So technical errors aside, intermediate values of methylation, show regions of high cell population heterogeneity. Here, we report a high level of cell population heterogeneity of methylation levels in CpG TL suggesting a novel flexible yet abundant mechanism of transcriptional regulation.

## Results

### CpG traffic lights determination

As has been shown many times, DNA methylation of a promoter can repress a corresponding gene. Nevertheless, correlation between gene expression and methylation of its promoter or body is not straightforward, suggesting the need to deconvolute DNA methylation profiles into the regions smaller than promoters. For this purpose, we focus on a methylation level of particular CpGs to investigate the link between methylation and expression. Following the logic previously reported in our works [40,47] where we used the reduced set of CpGs in the RRBS data, we expanded our previous approach and use whole-genome DNA methylation data (bisulfite sequencing, WGBS) and expression (RNA-seq) levels for 40 normal human primary cells and tissues from the Roadmap Epigenomics Project. We define CpG traffic lights as CpG dinucleotides with significant Spearman correlation coefficient (SCC) between DNA methylation and expression levels of a neighbouring gene (*FDR* < 0.1, Fig. 1).

**Figure 1.**
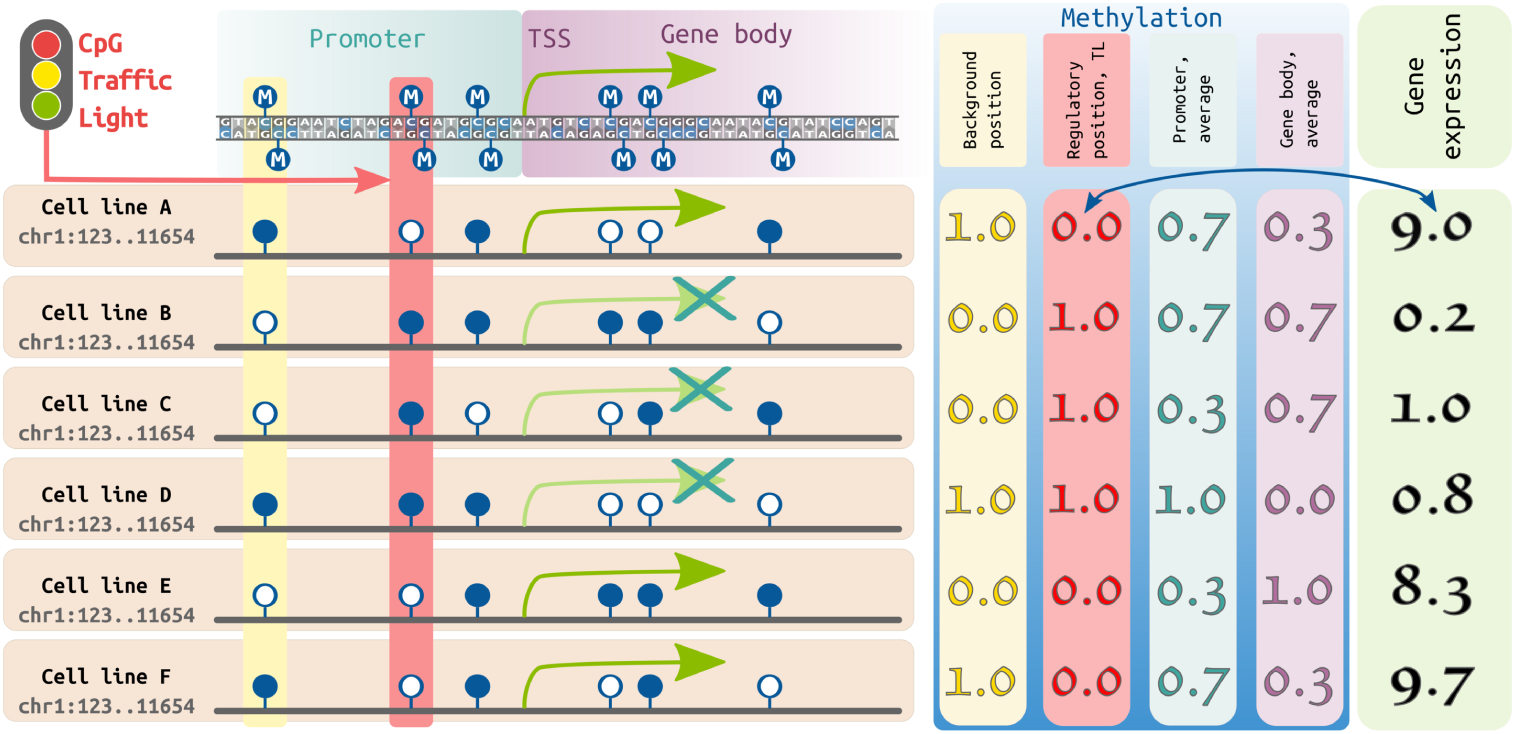
Schematic representation of a CpG traffic light determination. Left panel. Suppose we analyze a particular genomic region (chr1:123..11654), which contains for simplicity one gene, for 6 cell lines. For each CpG in this region and the gene we have methylation and expression vectors, respectively. CpG positions are represented by dark blue lollipops (filled: methylated CpG, empty: unmethylated CpG). First three CpGs are located within the promoter region, while the last three are located in gene body. Gene expression or lack of it is represented by green arrows. **Right panel.** A yellow column shows methylation of a random CpG (used as a background), methylation vector of this CpG demonstrates low correlation with gene expression (green box on the right, in RPKM). Correlation between an average promoter/gene body methylation (shown in light blue and light purple columns, respectively) and the corresponding gene expression is also low. However, for CpG TL (shown in red), methylation level significantly correlates with gene expression.

Here we state that the average methylation of promoter/gene body less frequently demonstrate a significant correlation with gene expression compared to the methylation of CpG TL, even then a proper multiplicity correction applied in each case. In particular, at the level of *FDR* < 0.1 we find only 44/58 genes for which average promoter/gene body methylation vectors correlate with expression vectors, while at the same level of significance we observe 6153 genes to correlate well with methylation levels of CpG TL. Other levels of significance demonstrate similar tendency (Table 1).

**Table 1.** Number of genes which have significant correlation between expression and methylation. Note: for multiplicity testing correction the number of genes was used in (1) and (2), while number of all CpG positions in each studied gene was used for the same purpose in (3). The (TTS) refers to a Transcript Termination Site.

| FDR-corrected p-value<br>(significance level) | Total number of genes, which have significant correlations between gene expression and methylation |  |  |
| --- | --- | --- | --- |
|  | average methylation of promoter regions (-1000..500) (1) | average methylation of gene bodies (+500..TTS) (2) | methylation of CpG TL (3) |
| 0.05 | 0 | 11 | 2706 |
| 0.1 | 44 | 58 | 6153 |
| 0.2 | 300 | 406 | 12,040 |

Among CpG TL, defined above (*FDR <* 0.1), the majority of those located in promoters demonstrate negative SCC, while the majority of those located in intron demonstrate positive SCC, and CpG TL in exons demonstrate similar number of both positive and negative SCC (Fig. 2). CpG TL are uniformly distributed along the genome (Manhattan plot, Supplementary Figure S1).

**Figure 2.**
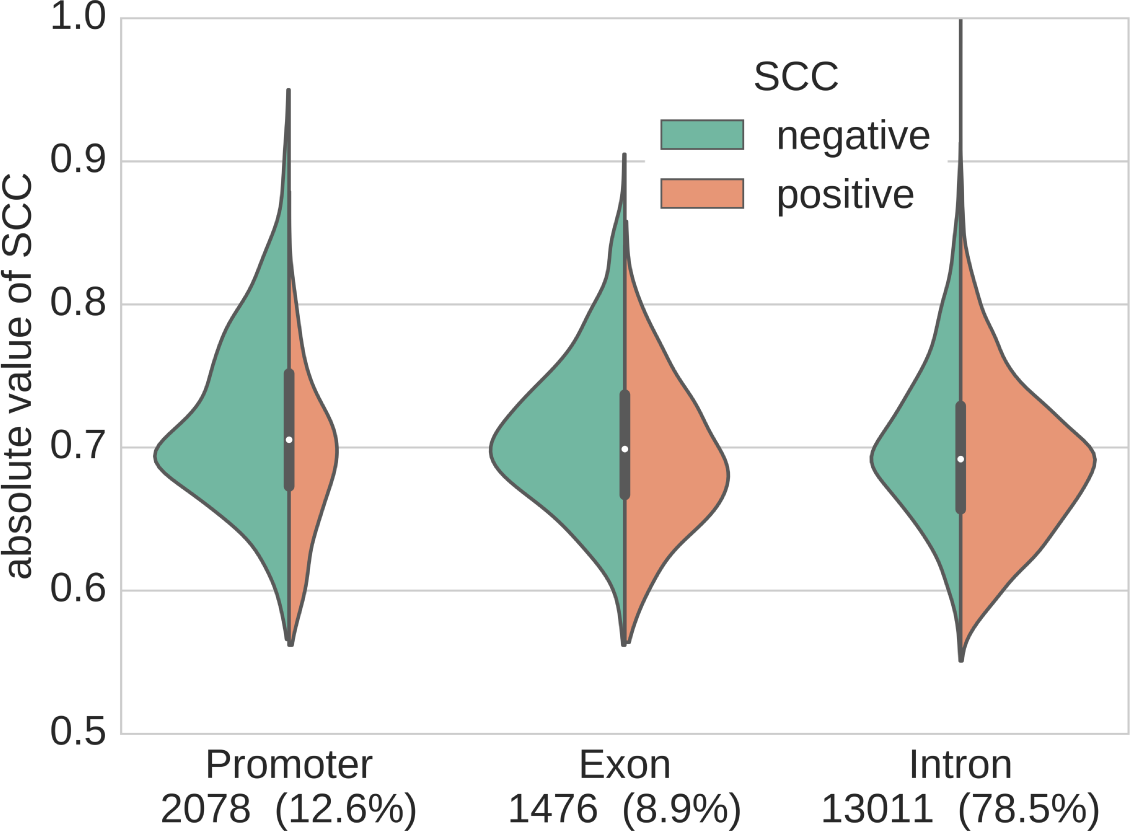
The distribution of SCC in the CpG TL. The total number of CpG TL in promoters, exons and introns are present at the bottom. Green (left) / pink (right) parts of the violin plots show the distribution of positive and negative SCC, respectively.

### CpG traffic lights are associated with highly heterogeneous genomic regions

A single CpG position can be either methylated or not, resulting in 0 or 1 methylation levels, in a diploid cell, allele specific methylation for some CpG positions can result in the methylation levels of 0.5. Allele-specific methylation has been reported to affect up to 10% of human genes [66] and is usually linked to genetic polymorphisms [58,66]. Since the allele specific methylation is usually linked to SNPs, intermediate methylation levels reported at the same genomic locations for several genetically unrelated samples represent heterogeneity of methylation levels among individual cells at a given CpG position. For the majority of CpG positions not detected as CpG TL (background CpGs, CpG BG, see Methods section for details), the levels of methylation were close either to 0 or 1 in all studied cell types (Fig. 3a,b), demonstrating homogeneity of the methylation levels in the cell population. At the same time the CpG TL with negative SCC between expression and methylation, both located in promoters and gene body, are intermediately methylated in many cell types (Fig. 3c,d). The similar tendency was observed for CpG TL with positive SCC (Supplementary Figure S2).

**Figure 3.**
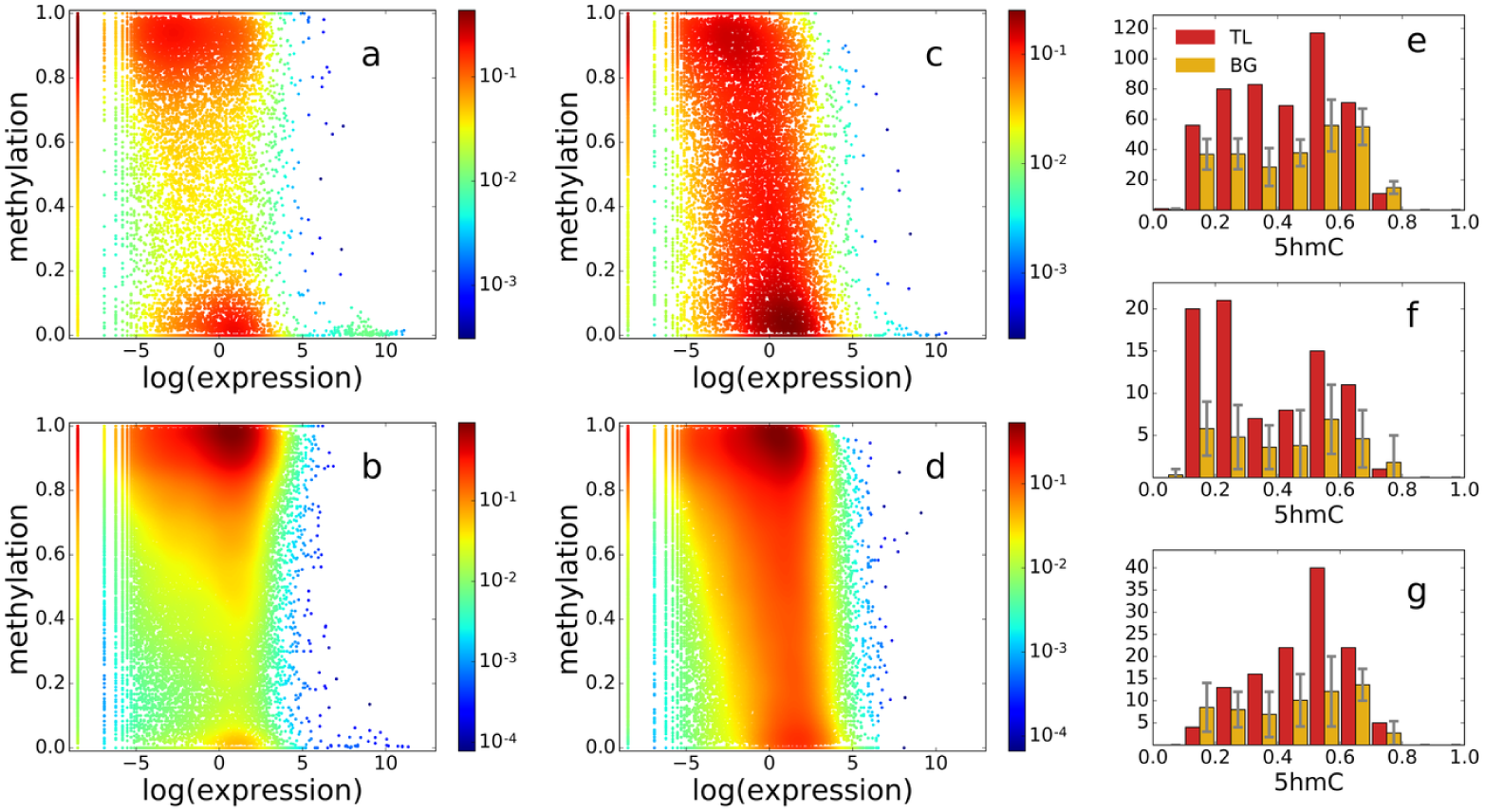
The distribution of CpG methylation and corresponding gene expression for CpG TL and background (negative SCC). The color represents the density of points in logarithmic scale. The distribution is shown for **(a)** random background, CpG BG in promoters (the number is equal to the number of CpG TL), **(b)** random background, CpG BG in gene bodies, **(c)** CpG TL in promoters (−1000…+500), **(d)** CpG TL in gene bodies (+500…TTS), **(e)** levels of 5hmC in CpG TL and CpG BG, **(f)** levels of 5hmC for CpG TL with positive and **(g)** negative causality score between DNA methylation and gene expression. Whiskers represent minimum/maximum out of the 10 random background samples

Since methylation levels of CpG TL are clearly more heterogeneous than that of background CpG dinucleotides, we decided to test whether methylation of these positions is also more dynamic in time. As a proxy of methylation dynamics we used levels of hydroxymethylcytosine (5hmC). Although the functional role of 5hmC is not fully elucidated, one of the most supported hypothesis is that 5hmC is an intermediate product of active DNA demethylation [17]. In standard bisulfite conversion experiments 5hmC cannot be distinguished from its precursor 5mC [24]. To compensate for that we used Illumina 450K oxBS-array data [16]. We report that CpG TL are enriched for 5hmC as compared to the CpG BG, supporting the idea of dynamic methylation in CpG TL (Fig 3e). This dynamic methylation of the CpG TL, caught by the snapshot experiment at different stages, might provide an explanation of the heterogeneity observed among them.

As a next step we divided CpG TL into subgroups based on causality scores, which allows one to computationally determine which of the vectors (methylation or expression) is a causal variable (see the Methods section for details). In our case, positive causality scores reflect cases where changes in DNA methylation cause the change in expression, whereas negative values of causality score correspond to CpG positions for which levels of methylation are a consequence of expression level. Surprisingly, it is mostly CpG TL with negative causality scores that demonstrate enrichment of high concentrations of 5hmC per site (Fig. 3g), which may suggest a positive feedback loop of the active transcription that activates DNA demethylation.

### CpG traffic lights are conserved across mammals and primates

To address functionality of CpG TL, we first investigate their evolutionary conservation. CpG TL are enriched with conserved positions both in mammals and in primates, estimated by GERP RS and PhyloP conservation scores, respectively (Fig. 4ab). Also, CpG TL are depleted in polymorphisms from ExAC (Fig 4c), as well as in repetitive sequences determined by both chromatin states (chromHMM, Fig. 4e) and repeatMasker (Fig 5a). This is in agreement with Eigen non-coding scores being significantly higher for CpG TL (Fig 4d). Moreover, gene enrichment analysis for GO terms shows that CpG TL are linked to genes involved in development, cell-to-cell communication and apoptosis (Table 2). Taken together, these results clearly suggest the functional role of CpG TL in the genome.

**Figure 4.**
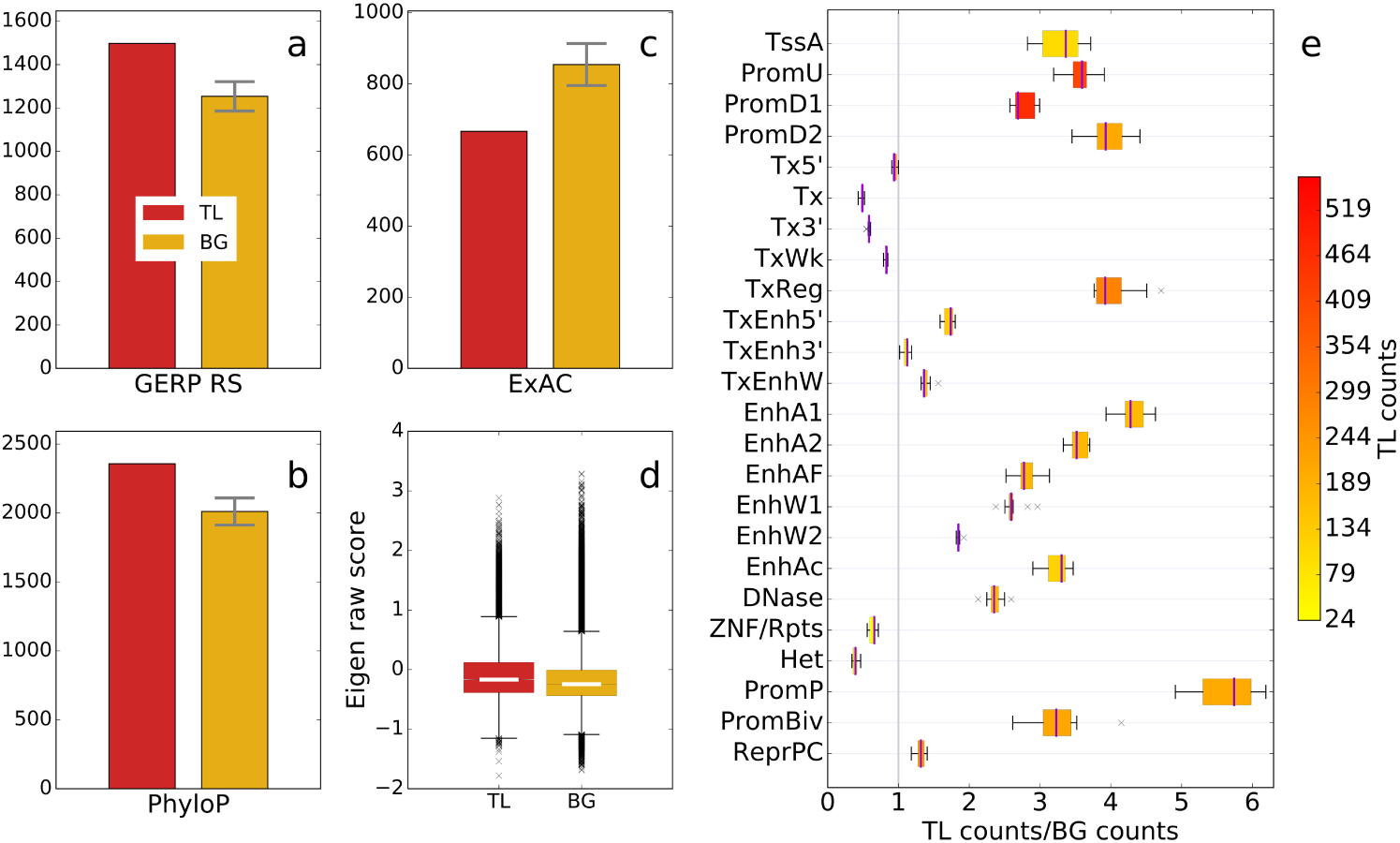
Number of CpG TL and BG sites demonstrating evolutionary conservation **(a)** in mammals and **(b)** in primates, **(c)** polymorphisms from ExAC, **(d)** Eigen non-coding functionality score, **(e)** averaged across 127 cell types ratio of TL / BG in chromatin states determined by chromHMM. Whiskers (abc) represent minimum / maximum out of the 10 random background samples. The color (e) reflects absolute number of CpG TL located in the given chromatin state.

**Figure 5.**
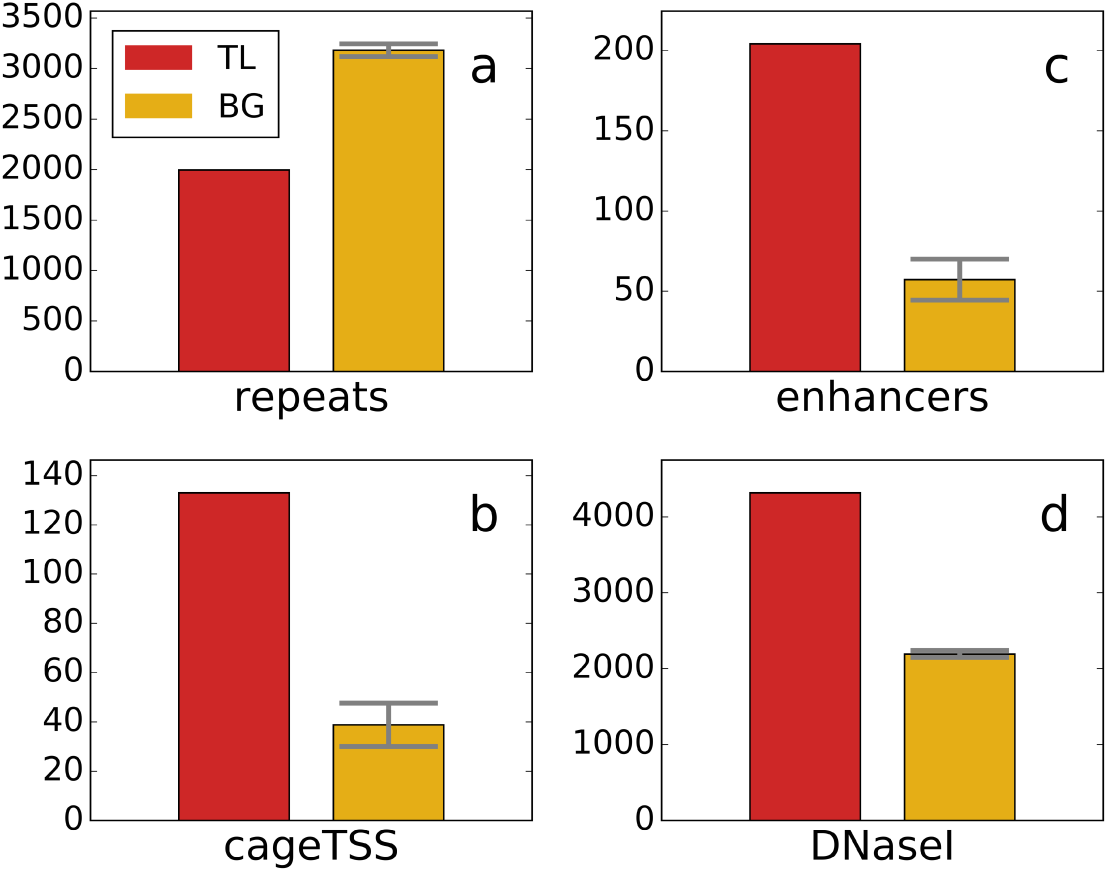
Frequency of CpG TL and CpG BG repeats **(a)**, TSS determined by CAGE **(b)**, enhancers **(c)** and DNase hypersensitive sites **(d)**. Whiskers represent minimum/maximum out of the 10 random background samples. All differences are significant (*p* – *value <* 5*E* – 5).

**Table 2.** Enrichment of CpG TL in biological processes (www.pantherdb.org)

| PANTHER<br>GO-Slim Biological<br>Process | Homo<br>sapi-<br>ens<br>(REF) | #<br>genes<br>with<br>CpG<br>TL | #<br>genes<br>ex-<br>pected | Fold En-<br>richment | +/- | P value |
| --- | --- | --- | --- | --- | --- | --- |
| developmental pro-<br>cess | 1938 | 627 | 451.14 | 1.39 | + | 2.11E-14 |
| cellular process | 8199 | 2160 | 1908.62 | 1.13 | + | 3.24E-11 |
| cell communication | 2674 | 787 | 622.47 | 1.26 | + | 1.21E-09 |
| cell adhesion | 481 | 190 | 111.97 | 1.7 | + | 1.57E-09 |
| biological adhesion | 481 | 190 | 111.97 | 1.7 | + | 1.57E-09 |
| system development | 1065 | 358 | 247.92 | 1.44 | + | 2.05E-09 |
| signal transduction | 2390 | 702 | 556.36 | 1.26 | + | 3.16E-08 |
| nervous system de-<br>velopment | 668 | 238 | 155.5 | 1.53 | + | 5.81E-08 |
| cell-cell adhesion | 305 | 124 | 71 | 1.75 | + | 1.43E-06 |
| intracellular signal<br>transduction | 991 | 312 | 230.69 | 1.35 | + | 2.46E-05 |
| mesoderm develop-<br>ment | 447 | 159 | 104.06 | 1.53 | + | 5.96E-05 |
| ectoderm develop-<br>ment | 405 | 142 | 94.28 | 1.51 | + | 5.32E-04 |
| heart development | 143 | 62 | 33.29 | 1.86 | + | 1.23E-03 |
| cellular component<br>movement | 413 | 137 | 96.14 | 1.42 | + | 1.03E-02 |
| mitosis | 372 | 122 | 86.6 | 1.41 | + | 4.05E-02 |
| induction of apopto-<br>sis | 85 | 38 | 19.79 | 1.92 | + | 4.12E-02 |
| transcription co-factors | 906 | 288 | 121.56 | 2.37 | + | 1.80E-46 |
| transcription fac-<br>tors | 1636 | 413 | 219.50 | 1.84 | + | 2.20E-38 |
| epigenetic regula-<br>tors | 692 | 221 | 92.84 | 2.38 | + | 4.48E-36 |
| histones | 72 | 5 | 9.66 | 0.52 | - | 0.48 |

### CpG traffic lights are enriched in transcription start sites, promoters and enhancers

To specify the functional role of CpG TL we tested various different genomic markups for the overrepresentation. We observe that CpG TL are enriched in all promoter types, determined by chromHMM [11], including active, bivalent and poised promoters (Fig. 4e). Interestingly, the strongest enrichment was observed in poised promoters (>3.5 times). Since the poised or bivalent chromatin is thought to be able to easily switch between active and repressed states [34], such enrichment may suggest a contribution of CpG TL to the maintenance of the bivalent state of the chromatin.

We also notice that CpG TL are highly enriched in all the chromatin states, corresponding to transcriptional start site (TSS). To dig deeper, we use TSS, determined by CAGE (Cap Analysis of Gene Expression), currently the most accurate technique to determine exact locations of TSS [12]. We determine 3.5-fold overrepresentation of CpG TL at the exact TSS position (Fig 5b). It should be noted that among all the groups of CpG TL located at TSS, the biggest group with the most pronounced overrepresentation over the background, has negative correlation and causality scores (Supplementary Figure S3). Negative causality score represents that changes in levels of expression drive the methylation levels, suggesting that for TSS regions methylation of a CpG TL is a marker, not the cause of expression.

Our data also show that CpG TL are enriched in various regulatory regions, yet the strongest enrichment is observed in enhancers, determined by CAGE bidirectional transcription (Fig 5a) and by chromatin states (Fig 4). Although all the enhancers are enriched for CpG TL, some types of enhancers are more prone to harbour them. We detect that among all enhancer categories the most enriched are various stem cell and hematopoietic cell enhancers (Fig. 6, Supplementary Table S1). All open chromatin regions determined as regions sensitive to DNaseI are also enriched for CpG TL (Fig 4e, 5d). On the other hand, as we reported before [40], CpG TL are not enriched in TFBS, if TFBS prediction is performed within the DNaseI sensitive regions.

**Figure 6.**
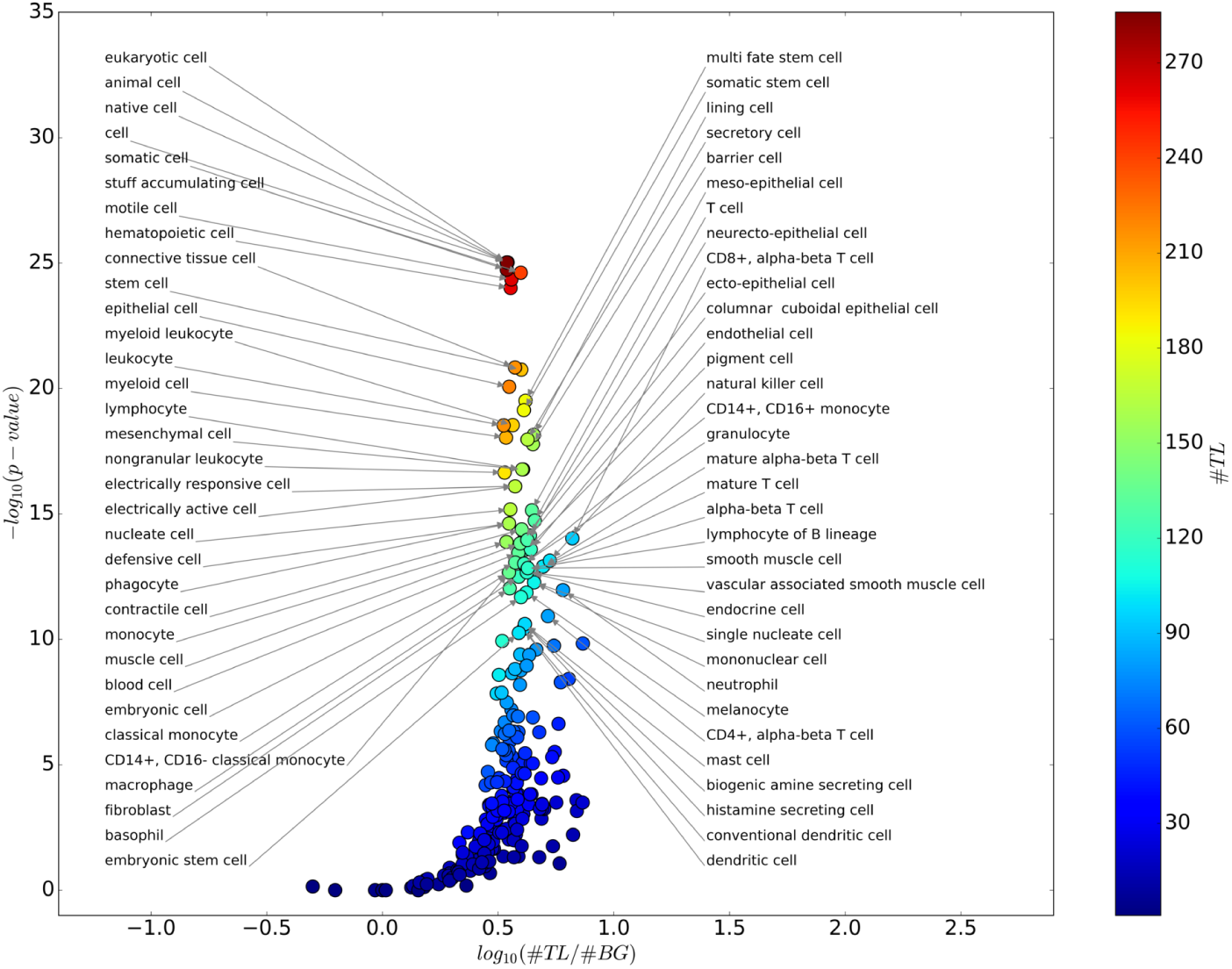
Functional categories of human enhancers enriched with CpG TL (negative SCC).

## Discussion

DNA methylation is tightly involved in regulation of gene expression in various normal and pathological processes. Therefore, this regulatory mechanism is an attractive target for therapies of the diseases with epigenetic abnormalities (reviewed in [52]). Modern technologies based on bisulfite sequencing allow for detection of DNA methylation with a single CpG dinucleotide resolution. Yet, at the stage of the downstream analysis methylation levels are averaged over the large regions. In this work we demonstrate that methylation profiles of particular single CpG dinucleotides (CpG TL) are more often significantly correlated with gene expression as compared to average promoter / gene body methylation even if for the multiplicity testing total number of CpG-gene pairs is used. It is a surprising observation, since it is widely accepted that DNA methyltransferases once bound to DNA move along [18] it or multimerize [59] methylating all neighbouring CpGs unless a boundary protein, such as Sp1, is bound in their way (reviewed in [63]). Yet, only a small fraction of CpG TL are located within the promoter and body of the same gene. We speculate that local change in DNA methylation can be achieved rather through active DNA demethylation, probably with the help of TET proteins, since byproduct of active demethylation, 5hmC is found to be overrepresented in CpG TL. However, a direct experiment, probably with the use of CRISPR/TALEN-based technology, is required to validate this hypothesis.

CpG TL are evolutionary conserved in both mammal and primate lineage, suggesting possible selection constraint, as well as depleted in SNPs, repeats and heterochromatin regions, supporting the hypothesis of CpG TL functionality. Genes that harbor CpG TL are associated with fundamental biological processes, such as development and signal transduction. CpG TL are also enriched in open chromatin and various regulatory regions, in particular at the exact TSS positions and active enhancers, especially those detected by bidirectional CAGE transcription [1]. This observation is in line with the recent reports that DNA methylatransferases DNMT3a/b are associated with enhancers and are important regulators of enhancer RNA production in hematopoietic stem cells [50]. Also, distal regulatory regions can initiate transcription themselves, being in turn regulated by DNA methylation [54], contributing to the similarity of TSS and enhcancers in terms of CpG TL enrichment.

In the light of overrepresentation in regulatory regions, depletion of CpG TL within TFBS is puzzling. One possible explanation would be that CpG TL are CpG dinucleotides located within the enhancers but outside the sites of regulatory protein binding. In this case, cytosine methylation accumulates as a consequence of the absence of TF binding [60,64], which makes methylation of CpG TL not a primary cause, but just a “passive” marker of absent gene expression resulting from inactivation of its regulatory element. Still, CpG traffic light methylation is a reliable marker of enhancer activity and gene expression, and can be used for practical applications.

Methylation levels of CpG TL vary between cell types dramatically and are characterized by increased frequency of intermediate methylation levels, indicating that only a fraction of cells within the same tissue have a certain CpG traffic light methylated. This variation cannot be attributed to genetic polymorphisms, since for our study we used samples from genetically different subjects, so it would be highly unlikely to have the same allele in a given position in the huge fraction of samples in the study. The more probable explanation is heterogeneity at the cell population level, which is indirectly supported by methylation dynamics in the form of increased levels of 5hmC. This heterogeneity is most likely stochastic, suggesting the role for CpG TL as a novel highly dynamic yet abundant markers of transcriptional regulation. This hypothesis is supported by the observation that the CpG TL are highly overrepresented at enhancers and poised promoters, suggesting their contribution into dynamics of expression.

## Conclusions

In this work we demonstrate that CpG TL are enriched in regulatory regions, including poised/bivalent promoters and enhancers.The mechanis of CpG traffic lights provide a promising insight into enhancer activity and gene regulation linking methylation of single CpG to expression.

## Methods

### DNA methylation and expression data processing

We selected 40 tissues and cell types (see Supplementary Table S2) for which both WGBS and RNA-seq data were available in Roadmap Epigenomics Project (NCBI). For WGBS data for each cell type we used three replicates with the highest number of reads and the best genome-mapping ratio. For 28 cell types 3 RNA-seq replicates were available, while for 7/5 cell types only 2/1 replicate were available respectively. All WGBS data and the majority (95 out of 103) RNA-seq files were obtained by Illumina, while 8 RNA-seq files were obtained by SOLiD. The quality of all files were checked with FastQC [2]. For all files sequenced by Illumina read trimming and adapter removal were performed by Trimmomatic [6] (adapters available at Epigenomics Project (NCBI); up to 2 mismatches between an adapter and a read sequence; 5bp sliding window; quality threshold of 20; removing sequences if their length after trimming becomes less than 20 bp) For the SOLiD samples we used Cutadapt [37] (adapters from NCBI, up to 10% error rate relative to the length of the matching region; quality threshold of 20; removing sequences if their length after trimming becomes less than 20 bp).

We mapped WGBS data to the genome (assembly GRCh38-Ensembl 78) with Bismark [30] (zero mismatches permitted in the seed, 20bp seed length, 0/500bp the min/max insert size for valid paired-end alignments). We used only methylated cytosines in CpG context, covered with not less than 4 reads. For each CpG position in each of the 40 samples, the methylation values were averaged from the three replicates per sample. We removed a CpG position if methylation values were available for less than 20 samples. We also removed a CpG position if methylation values were the same value for all the samples (i.e. all zeros, all ones, etc.).

We mapped RNA-seq data to the genome (assembly GRCh38-Ensembl 78) with Tophat v2.0.13 [62] (allowing for up to 2 mismatches and 2 gaps per read, reporting read alignments for paired-end reads only if both reads in a pair can be mapped). We generated expression matrix using the FeatureCount [35], while the expression profiles were normalized to RPKM values. Genes with zero reads in all samples were excluded. The expression profiles were normalized to a range [0, 1] to match the range with the one of the methylation profile:

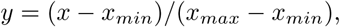

where *y* is normalized expression value for the gene, *x* is RPKM value for the gene, *x_min_* and *x_max_* are minimum and maximum expression (RPKM) values respectively.

### CpG traffic lights detection

To determine CpG TL we considered all pairs of genes and CpGs located within 1000 bp upstream of TSS to 3’ gene end (genome assembly GRCh38-Ensembl 78). One CpG might be associated with multiple genes, similarly, one gene might be associated with multiple CpGs. For each CpG-gene pair we created two 20-40-dimensional vectors of methylation levels [0, 1] and normalized gene expression [0, 1], we further refer to each of the two vectors as a methylation and expression profiles. In total we had 1,774,602 CpGs associated with 46,692 genes (which gives 1,963,205 pairs).

For each CpG position, we calculated SCC between the methylation and expression profiles for all available samples. FDR was performed by Benjamini-Hochberg procedure for correction for multiplicity testing for the total number of position-gene pairs. We called a CpG position a CpG traffic light (CpG TL) if it had a significant correlation coefficient between methylation and expression profiles at the level of *FDR* < 0.1 (unless explicitly mentioned otherwise). We found 16,178 such CpG TL (0.9% of the original number of CpGs) that correspond to 6153 genes.

We also calculated a causality score between methylation and expression profiles to computationally assess the pairwise causal direction between these two variables. We used a pairwise linear non-Gaussian acyclic model, LINGAM [20] to calculate the likelihood ratio defined as follows:

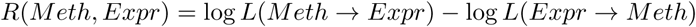

The positive causality means that the change in methylation is expected to cause the expression change, and vice versa for the negative causality values: expression determines methylation. It should be noted that the range for possible causality scores depends on the number of samples. Since for different CpG positions we used various numbers of samples (20-40), we normalized causality scores to the normal distribution N(0, 1). To make the causality scores directly comparable between CpG positions, we performed this normalization independently for each group of CpGs that have the same profile length. To avoid noise in the causality scores, we did not consider values close to 0 (between −1 and +1) and for simplicity we call “positive” the values that are higher than 1 and “negative” the values that were smaller than −1.

### Construction of background datasets

We aimed to explore enrichment with CpG TL inside various genomic regions. For this purpose we needed to have an equal size background set. For every CpG TL position we selected a random background CpG position (CpG BG) with not more than 5% difference for both GC- and CpG contents in 200bp window, as some genomic annotations are sensitive to GC- and CpG- content. We repeated the selection process 10 times to obtain 10 different independent background sets.

For heatmaps (Fig. 2) we selected CpG TL with negative SCC and an equal size random background set, split all the CpGs into promoter regions [*TSS*–1000, *TSS*+500] and gene body [*TSS* + 500, end of the gene] and created density plots using gaussia_kdefrom scipy.stats.

Recently, we have shown that methylated CpGs within CpG islands are more preserved from substitutions in primate evolution even in the same local GC- and CpG content [46]. Yet, the algorithms for CpG islands search use some arbitrary parameters and may not be accurate in determination boundaries [38]. Therefore, controlling for a presence of a CpG island would not necessarily reduce the bias. However, CpG islands usually contain internal sequence patterns, for example transcriptional factor binding sites [39]. So, we tested if regions of a 100 bp around randomly selected 1000 CpG TL (CpG TL excluded) contain different sequence motifs that the one of the CpG BG [31]. The obtained motifs were fairly similar for CpG TL and CpG BG sequences (Supplementary Figure S4), suggesting that control for GC- and CpG contexts is sufficient for the purpose of this study.

### Genomic annotations

We annotated all CpG positions with overlapping genomic features. For each feature we calculated the number of CpG TL and background positions located within the annotation. To test the significance of the overrepresentation we used the exact Fisher test. Additionally, we calculated the overrepresentation for CpG TL with positive/negative SCC/causality scores separately.

To address the frequency of 5-hydroxymethylcytosine in CpG TL we used oxidative-bisulfite (oxBS) assay data from human cerebellum (GEO, GSE63179) [16]. We converted the coordinates to genomic ranges with the help of R Bioconductor ‘minfi’ [3] package and to hg38 with liftOver. Four oxBS replicates were averaged.

We use repeats obtained by RepeatMasker for hg38 track from USCS Genome Browser hgdownload.soe.ucsc.edu/goldenPath/hg38/database/rmsk.txt.gz

We obtained the robust CAGE clusters [12] from fantom.gsc.riken.jp/5/data/ and the robust hg19 enhancers [1] from FANTOM5 from (enhancer.binf.ku.dk/presets/robust_enhancers.bed) and mapped them to hg38 with the liftOver.

The DNaseI hypersensitivity clusters were downloaded from UCSC Genome Browser (hgdownload.soe.ucsc.edu/goldenPath/hg38/database/wgEncodeRegDnaseClustered.txt.gz)

#### Conservation and Eigen scores

Conservation of CpG TL and background sites in mammalian and primate lineages was assessed with UCSC Genome Browser GERP RS [8] and PhyloP [49] hg19 tracks, respectively. We calculated how many sites in each dataset have GERP RS score greater than 2, which we considered as conserved in mammals and PhyloP score greater than 0.5, which we considered conserved in primates. Overall functional scores for each site were calculated with Eigen, an approach to predict functionality of non-coding variants using different annotations [21]. Higher Eigen scores imply more likely functionality of respective genome sites.

#### TFBS

For transcriptional factor binding site (TFBS) prediction we used models provided in HOCOMOCO v10 [32]. PWM thresholds were selected according to the pre-calculated the *P* – *value* < 0.0005 (i.e., when 5 of 10,000 random words had scores no less than the thresholds). Out of all predicted TFBS we considered only those present in DNaseI hypersensitivity regions.

#### ChromHMM

The Roadmap Epigenomics Consortium 25-state segmentation of 127 epigenomes predicted with ChromHMM [11,51] was used to assess chromatin location of CpG TL. The annotation in based on the imputed data for 12 chromatin marks (H3K4me1, H3K4me2, H3K4me3, H3K9ac, H3K27ac, H4K20me1, H3K79me2, H3K36me3, H3K9me3, H3K27me3, H2A.Z, and DNaseI). The annotations were downloaded from egg2.wustl.edu/roadmap/web_portal/imputed.html#chr_imp.

Each of the CpG TL/background datasets was characterized by an average frequency of a CpG from a dataset to be located in one of the 25 chromatin states.

#### Gene enrichment analysis

For the gene ontology enrichment analysis (GO) we used Panther [43] with default parameters and Bonferroni correction. Separately, we tested for the enrichment of genes with CpG TL among transcriptional factors and co-factors [55] and epigenetic regulators [41] using Fisher exact test with Bonferroni correction.

## Authors contribution

AK processed the raw data and contributed to data analysis; AL contributed to data processing and performed the overrepresentation analysis; VR performed data analysis; AA contributed to statistical analysis; VBB contributed to study design; YAM designed the study and drafted the MS. All authors contributed to MS preparation.

## Acknowledgments

This work was supported by RFBR grant 14-04-00180 to YAM. VBB is supported by the base research fund of the King Abdullah University of Science and Technology (KAUST).

## Supplementary materials

**Supplementary Figure S1.**
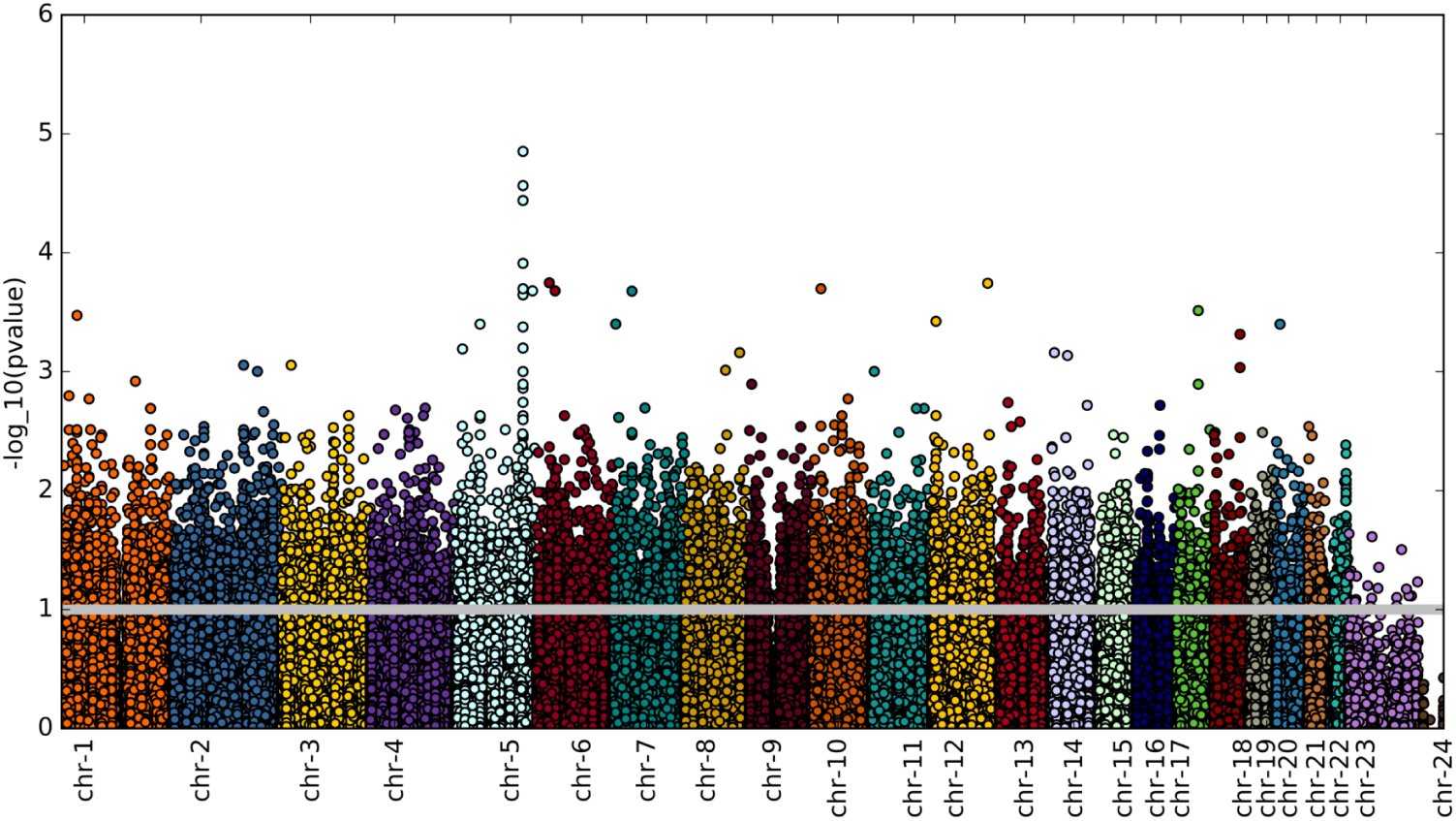
Distribution of CpG TLs along the genome.

**Supplementary Figure S2.**
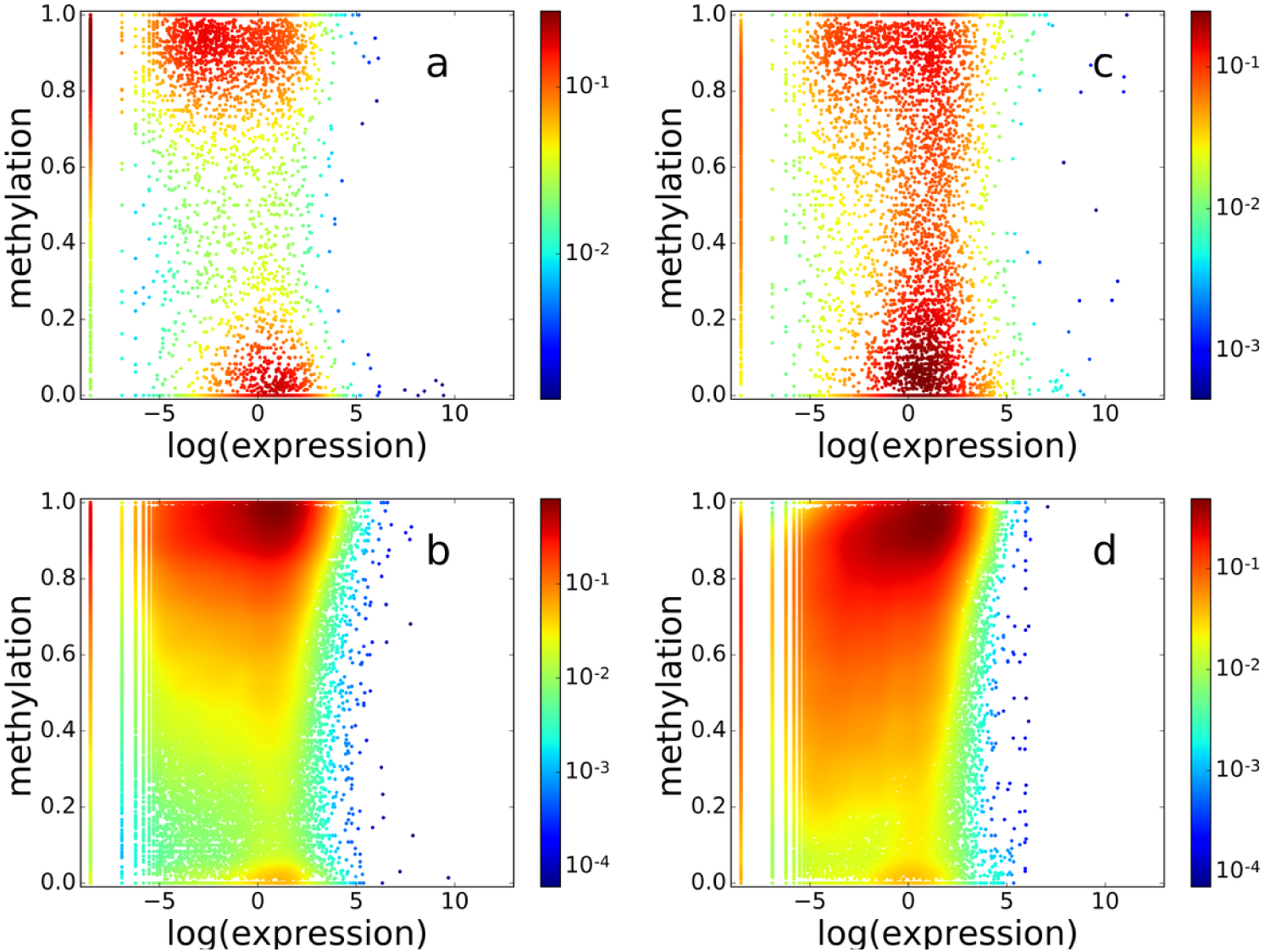
The distribution of CpG methylation and corresponding gene expression for CpG TLs and background (positive SCC). The color represents the density of points in logarithmic scale. The distribution is shown for **(a)** random background CpG (BG) in promoters (the size is equal to the number of CpG TL points), **(b)** random equal size BG in gene bodies, **(c)** CpG TL in promoters (−1000…+500), **(d)** CpG TL in gene bodies (+500…TTS)

**Supplementary Figure S3.**
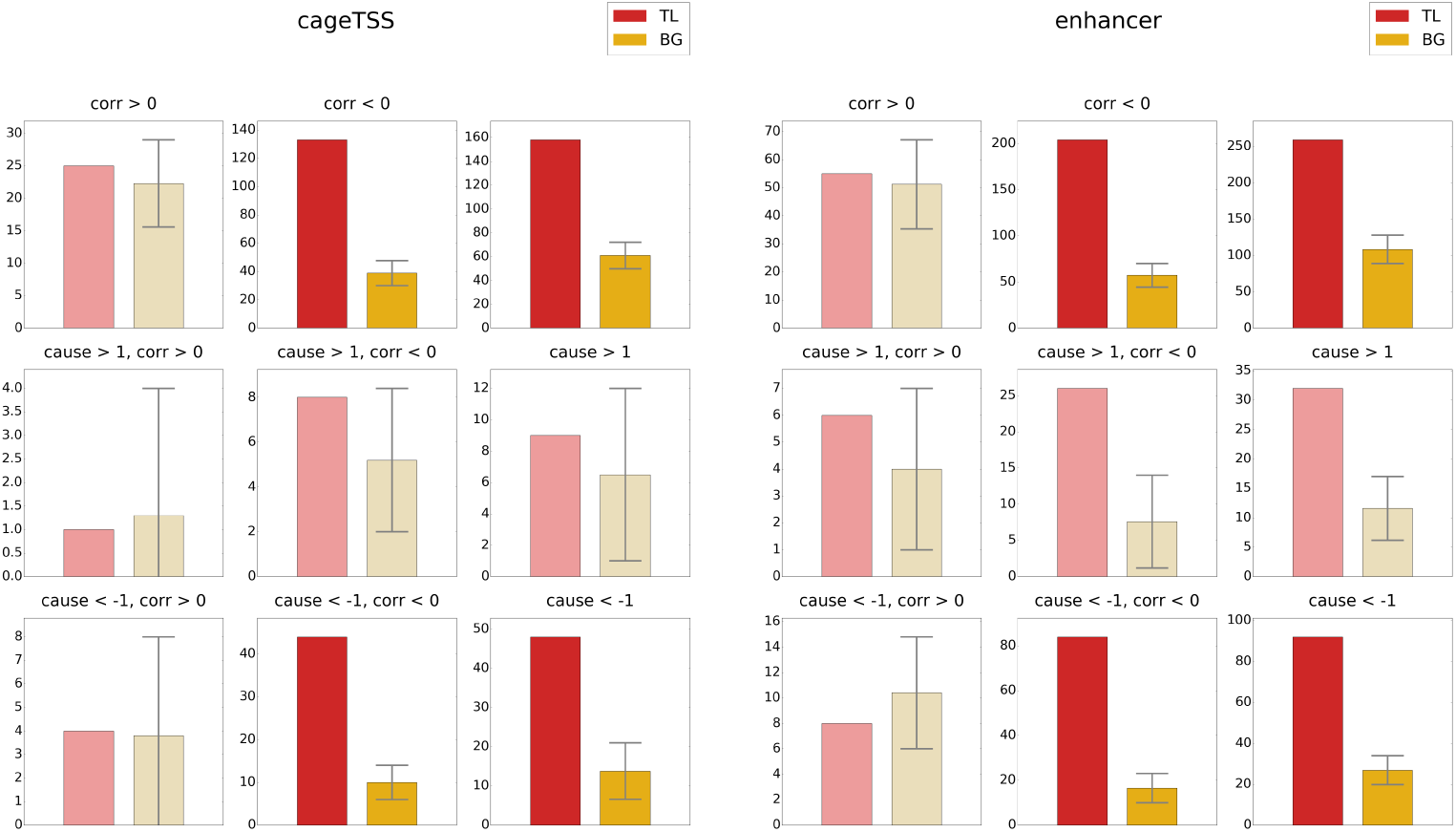
Frequencies of CpG TL and BG in CAGE TSS and enhancers. CpG TL are separated in groups based on the sign of correlation and causality score.

**Supplementary Figure S4.**
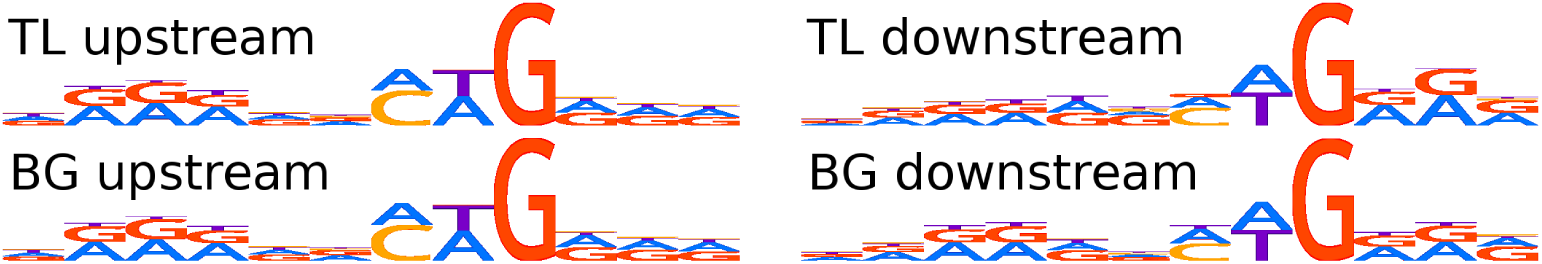
Motives near CpG.

**Supplementary Table S1.** Most enriched with CpG TLs categories of enhancers

| Enhancer category | CpG TL/BG ratio | P value |
| --- | --- | --- |
| CD8+, alpha-beta T cell | 6.620 | 9.475E-15 |
| neutrophil | 6.029 | 1.090E-12 |
| natural killer cell | 5.294 | 7.295E-14 |
| CD4+, alpha-beta T cell | 5.190 | 1.187E-11 |
| mature alpha-beta T cell | 4.951 | 1.251E-13 |
| mature T cell | 4.951 | 1.251E-13 |
| alpha-beta T cell | 4.951 | 1.251E-13 |
| T cell | 4.555 | 1.866E-15 |
| single nucleate cell | 4.526 | 5.416E-13 |
| mononuclear cell | 4.526 | 5.416E-13 |
| lining cell | 4.511 | 7.098E-19 |
| barrier cell | 4.473 | 1.659E-18 |
| meso-epithelial cell | 4.430 | 7.258E-16 |
| endothelial cell | 4.380 | 2.606E-14 |
| endocrine cell | 4.346 | 2.165E-13 |
| neurecto-epithelial cell | 4.340 | 7.265E-15 |
| lymphocyte of B lineage | 4.249 | 1.453E-13 |
| secretory cell | 4.238 | 1.083E-18 |
| columnar cuboidal epithelial cell | 4.228 | 1.090E-14 |
| vascular associated smooth muscle cell | 4.216 | 2.165E-13 |
| melanocyte | 4.192 | 1.376E-12 |
| pigment cell | 4.189 | 2.606E-14 |
| smooth muscle cell | 4.173 | 1.453E-13 |
| multi fate stem cell | 4.153 | 3.080E-20 |
| mast cell | 4.128 | 2.471E-11 |
| biogenic amine secreting cell | 4.128 | 2.471E-11 |
| histamine secreting cell | 4.128 | 2.471E-11 |
| CD14+, CD16+ monocyte | 4.102 | 9.677E-14 |
| somatic stem cell | 4.089 | 7.315E-20 |
| ecto-epithelial cell | 4.087 | 1.080E-14 |
| granulocyte | 4.073 | 9.677E-14 |
| conventional dendritic cell | 4.068 | 3.896E-11 |
| dendritic cell | 4.068 | 3.896E-11 |
| mesenchymal cell | 4.061 | 1.692E-17 |
| lymphocyte | 4.020 | 1.692E-17 |
| contractile cell | 4.000 | 4.174E-15 |
| stem cell | 3.984 | 1.781E-21 |
| basophil | 3.972 | 2.072E-12 |
| stuff accumulating cell | 3.964 | 2.444E-25 |
| muscle cell | 3.936 | 1.502E-14 |
| embryonic stem cell | 3.885 | 5.567E-11 |
| macrophage | 3.883 | 3.182E-13 |
| blood cell | 3.866 | 3.574E-14 |
| embryonic cell | 3.754 | 8.707E-14 |
| electrically responsive cell | 3.747 | 7.984E-17 |
| electrically active cell | 3.747 | 7.984E-17 |
| connective tissue cell | 3.744 | 1.465E-21 |
| myeloid leukocyte | 3.660 | 2.904E-19 |
| motile cell | 3.609 | 4.564E-25 |
| hematopoietic cell | 3.581 | 9.966E-25 |
| nucleate cell | 3.571 | 6.678E-16 |
| fibroblast | 3.547 | 9.294E-13 |
| epithelial cell | 3.530 | 8.584E-21 |
| defensive cell | 3.524 | 2.416E-15 |
| phagocyte | 3.524 | 2.416E-15 |
| classical monocyte | 3.523 | 2.165E-13 |
| CD14+, CD16- classical monocyte | 3.523 | 2.165E-13 |
| cell | 3.476 | 9.456E-26 |
| eukaryotic cell | 3.454 | 9.456E-26 |
| animal cell | 3.454 | 9.456E-26 |
| somatic cell | 3.451 | 1.852E-25 |
| native cell | 3.446 | 9.456E-26 |
| monocyte | 3.438 | 1.313E-14 |
| myeloid cell | 3.422 | 9.128E-19 |
| nongranular leukocyte | 3.375 | 2.252E-17 |
| leukocyte | 3.344 | 2.965E-19 |

**Table S2.** Names of the cell samples in the study

| Sample Name | Number of Methylation Replicates | Number of Expression Replicates |
| --- | --- | --- |
| bladder | 3 | 1 |
| CD14 primary cells | 3 | 1 |
| CD56 primary cells | 3 | 1 |
| H9 cell line | 3 | 1 |
| iPS DF 6.9 cell line | 3 | 1 |
| CD3 primary cells | 3 | 2 |
| H1 +BMP4 cell line | 3 | 2 |
| H1 BMP4 derived mesendoderm cultured cells | 3 | 2 |
| H1 derived mesenchymal stem cells | 3 | 2 |
| iPS DF 19.11 cell line | 3 | 2 |
| liver | 3 | 2 |
| placenta | 3 | 2 |
| adipose | 3 | 3 |
| adrenal gland | 3 | 3 |
| esophagus | 3 | 3 |
| gastric | 3 | 3 |
| H1 cell line | 3 | 3 |
| H1 derived neuronal progenitor cultured cells | 3 | 3 |
| heart aorta | 3 | 3 |
| heart left ventricle | 3 | 3 |
| heart right atrium | 3 | 3 |
| heart right ventricle | 3 | 3 |
| hESC-derived CD184+ endoderm cultured cells | 3 | 3 |
| hESC-derived CD56+ ectoderm cultured cells | 3 | 3 |
| hESC-derived CD56+ mesoderm cultured cells | 3 | 3 |
| HUES64 cell line | 3 | 3 |
| IMR90 cell line | 3 | 3 |
| large intestine | 3 | 3 |
| lung | 3 | 3 |
| muscle leg | 3 | 3 |
| muscle trunk | 3 | 3 |
| ovary | 3 | 3 |
| pancreas | 3 | 3 |
| psoas muscle | 3 | 3 |
| sigmoid colon | 3 | 3 |
| small intestine | 3 | 3 |
| spinal cord | 3 | 3 |
| spleen | 3 | 3 |
| stomach | 3 | 3 |
| thymus | 3 | 3 |

